# Land Use Change Increases Wildlife Parasite Diversity in Anamalai Hills, Western Ghats, India

**DOI:** 10.1101/645044

**Authors:** Debapriyo Chakraborty, D. Mahender Reddy, Sunil Tiwari, Govindhaswamy Umapathy

**Affiliations:** CSIR-Laboratory for the Conservation of Endangered Species, Centre for Cellular and Molecular Biology, Hyderabad 500048, India; EP57 P C Ghosh Road, Kolkata 700048, India

**Keywords:** Parasite diversity, Land use change, Plantation, Livestock, Human settlement, Anamalai hills, Habitat fragmentation, Rainforest

## Abstract

Anthropogenic landscape change such as land use change and habitat fragmentation are known to alter wildlife diversity. Since host and parasite diversities are strongly connected, landscape changes are also likely to change wildlife parasite diversity with implication for wildlife health. However, research linking anthropogenic landscape change and wildlife parasite diversity is limited, especially comparing effects of land use change and habitat fragmentation, which often cooccur but may affect parasite diversity substantially differently. Here, we assessed how anthropogenic land use change (presence of plantation, livestock foraging and human settlement) and habitat fragmentation may change the gastrointestinal parasite diversity of wild mammalian host species (n=23) in Anamalai hills, India. We found that presence of plantations, and potentially livestock, significantly increased parasite diversity due possibly to spillover of parasites from livestock to wildlife. However, effect of habitat fragmentation on parasite diversity was not significant. Together, our results showed how human activities may increase wildlife parasite diversity within human-dominated landscape and highlighted the complex pattern of parasite diversity distribution as a result of cooccurrence of multiple anthropogenic landscape changes.

## INTRODUCTION

Land use change and habitat fragmentation are two major landscape-level outcomes of human activities that significantly impact biodiversity ^1–3^. Consequently, considerable research on biodiversity change in human-dominated landscape have been conducted, which has resulted in improved understanding of how these two human impacts on landscape can impact biodiversity ^1,4,5^. These anthropogenic factors can also modify host– parasite interactions, which, in turn, can lead to either increase or decrease in parasite diversity ^6–8,8^. Understanding how these factors may influence parasite diversity is ecologically important for multiple reasons. For instance, parasites regulate host population dynamics ^9^, alter species communities ^10^ and constitute a significant proportion of total biomass of any ecosystem ^11^, which is not surprising considering parasites comprise at least 40% of all animal species on earth ^12^. Despite their ecological importance, our knowledge on parasite diversity is limited^13,14^, particularly in the context of increasing human impact on environment, underlining a significant research gap^15,16^. The gap is specifically wide for wildlife hosts and urgent research is required in the face of recent increased emergence of novel pathogens of wildlife origin ^7,17,18^. It is, thus, crucial to answer how anthropogenic land use change and habitat fragmentation may impact parasite diversity in the wild.

Land use change can affect parasites both directly and indirectly. By altering environment (for example, through pollution), land use change may render transmission of environmentally-transmitted parasites difficult. This is particularly true for parasites that has life stages outside host body. However, land use change can indirectly impact parasite diversity by altering host diversity as it is one of the strongest predictors of parasite diversity ^19–22^. By decreasing host diversity and abundance, land use change can deplete richness of parasites particularly those that require multiple obligatory hosts ^23^. This is evident when many host species that are threatened in their natural habitat appear to harbour fewer parasites ^24^. On the other hand, land use change can also increase parasite diversity in multiple ways. Land use change can increase parasite diversity by increasing host diversity. For instance, land use change such as agricultural field or land-fill can act as resource traps and amplify host diversity artificially ^25^. Land use change can also increase parasite diversity by introducing non-native parasites such as parasites of domestic and feral animals and even from humans ^26^.

It is also important to distinguish between different types of land use change and their effects on parasites ^27^. One type of land use change that has not been studied well is the effect of plantation on wildlife parasites ^27^. Plantations are usually monocultures of exotic or native plant species grown as timber or fuel wood or as cash crops and have a large and increasing footprint in wildlife habitats worldwide ^28^. They can sometime act as refuge to wildlife but usually with a biotic homogenising effect^29,30^. Consequently, plantation may also increase but homogenise parasite community. Plantations are often accompanied by settlement of labourers and livestock foraging ^31,32^. These changes within a wildlife habitat can both increase or decrease parasite diversity. Parasite diversity may decrease if wildlife hosts avoid human areas to lessen confrontation with humans and resource competition with livestock. On the other hand, generalist species may actually thrive in human settlements by utilizing novel resources ^33,34^. Herbivore species may also prefer to stay closer to human settlements and livestock (“spatial refugia”) that may displace predators ^35–38^. Moreover, many wildlife, over time, may actually get habituated to humans and livestock and aggregate near human-dominated landscape ^39,40^. These aggregations may eventually increase parasite diversity by increasing contact between native host species. Such situations may also increasingly expose wildlife to humans and human-associated animals such as livestock and commensals, increasing chance of spillover of non-native parasites to wildlife.

Habitat fragmentation may lead to higher parasite diversity because heavily fragmented habitats may disrupt wildlife dispersal and increase host diversity in smaller fragments. Such increase in host diversity in a smaller patch may alter host characteristics such as home range, abundance and intra and interspecific contacts thus increasing overlap among host species making host individuals exposed to higher parasite infections ^41,42^. These effects are likely to be greatest in the smallest and most isolated of the fragments ^3,43^. By disrupting host dispersal, fragmentations can also adversely affect parasite diversity. This could be especially true for parasites who require multiple host species to complete its life cycle, such as those that are transmitted trophically ^44^. So far, many studies looked into this effect but the results have been mixed ^6,41,45–47^.

The Anamalai (Elephant hills in *Tamil*) hills of southern India is a highly biodiverse rainforest habitat of Western Ghats, which holds about 30% of India’s plant and vertebrate species diversity in less than 6% of the country’s area ^48^. It is also one of the most altered natural habitats in India and typifies different levels of land use change and habitat fragmentation rampant in Indian wildlife habitats. Large section of the habitat is highly modified due to land use change, bordered by large, relatively undisturbed tropical rainforests. The landscape is a matrix of over 40 rainforest fragments (1-2,500 ha in size),often surrounded by plantations (coffee, tea and cardamom), roads, hydroelectric dams and settlements ^49^. Highly-modified fragments contain within them human settlements and have higher livestock pressures than other remote, less disturbed fragments. In spite of such high levels of land use change and habitat fragmentation, the Anamalai hills still harbour a large number of wildlife whose ranges often unavoidably overlap with humans and livestock ^50–53^. In fact, large number of wildlife species are regularly observed within human-dominated habitats and this concurrence with humans often precipitates into wildlife-human conflicts ^49,50,54–56^. It is possible that many of the wildlife are important reservoirs of multiple environmentally-transmitted parasites. In fact, recent studies have recorded important parasite groups within certain host species populations that may cause Ascariasis, Trichuriasis and Strongylodiasis in humans ^45,46,57,58^.

To assess the effect of land use change (plantation, livestock foraging and human settlements) and habitat fragmentation on parasite diversity, we studied gastrointestinal parasites of wild mammalian hosts across rainforest fragments in Anamalai hills. Using statistical models, we tested effects of land use change and habitat fragmentation on parasite diversity. We predicted a positive impact of land use change on parasite diversity due to increased host diversity and an increased exposure of wildlife to humans and livestock. For habitat fragmentation too, we predicted an increase in parasite diversity with decrease in habitat size and increase in habitat isolation. Our alternative predictions were that land use change and habitat fragmentation could actually deplete parasite diversity by decreasing host diversity in disturbed fragments. Finally, land use change and habitat fragmentation may not significantly impact parasite diversity either by not impacting host community or by not spillover from non-native hosts such as livestock and humans.

## MATERIALS AND METHODS

### Ethical statement

For this study, faecal samples were collected only noninvasively. As a result, no animal was sacrificed or harmed during sampling. Part of the sampling was done within Anamalai Tiger Reserve, which is a protected area. Hence, appropriate written permission was taken from the Tamil Nadu Forest Department (Letter Ref. No. WL 5/58890/2008, dated 2nd September 2009).

### Study site

Located south of the Palghat gap (11° N) of the Western Ghats, Anamalai hillss once had large tracts of tropical rainforest dotted with few tribal settlements. Between 1860 and 1930, British colonisers started clearing the rainforests extensively for cultivation of tea and coffee and developing teak and *Eucalyptus* plantations, particularly in the Valparai Plateau ^59^. As a result, the Anamalais today consists of both a relatively undisturbed, large (958.59 km^2^) tropical rainforest within the protected Anamalai Tiger Reserve (ATR; 10°12′–10°35′N and 76°49′–77°24′E) and about 1,000 ha highly degraded Valparai Plateau (Figure 1). The plateau consists of many tea estates and other plantations, which are surrounded by four protected areas—ATR in Tamil Nadu state and three others in Kerala state. The major vegetation types include scrub forests in the rain-shadow areas in the eastern foothills, dry and moist deciduous forests (<800 m), mid-elevation tropical wet evergreen forest (600-1,500 m) and high-altitude shola-grassland ecosystems (>1,500 m) ^60^. Although a large part of the tropical wet evergreen forests occurs within ATR, many of the smaller (< 200 ha) fragments are found in private estates in the Valparai plateau. These small fragments are highly degraded and disturbed due to fuel-wood collection and livestock grazing. Valparai town also is a part of the plateau and around 200,000 people live across the town and plantations ^60^. Due to the ongoing habitat fragmentation, the whole landscape is a matrix of over 40 rainforest fragments, ranging 1 ha-2,500 ha in size and often surrounded by plantations (coffee, tea and cardamom), roads, hydroelectric dams and settlements ^49^. Based on size range (2-2,500 ha), level of perceived human disturbance and access, we selected 19 mid-elevation tropical rainforest and three low-elevation dry and moist deciduous forest fragments for sampling (Figure 1).

**Figure 1.**
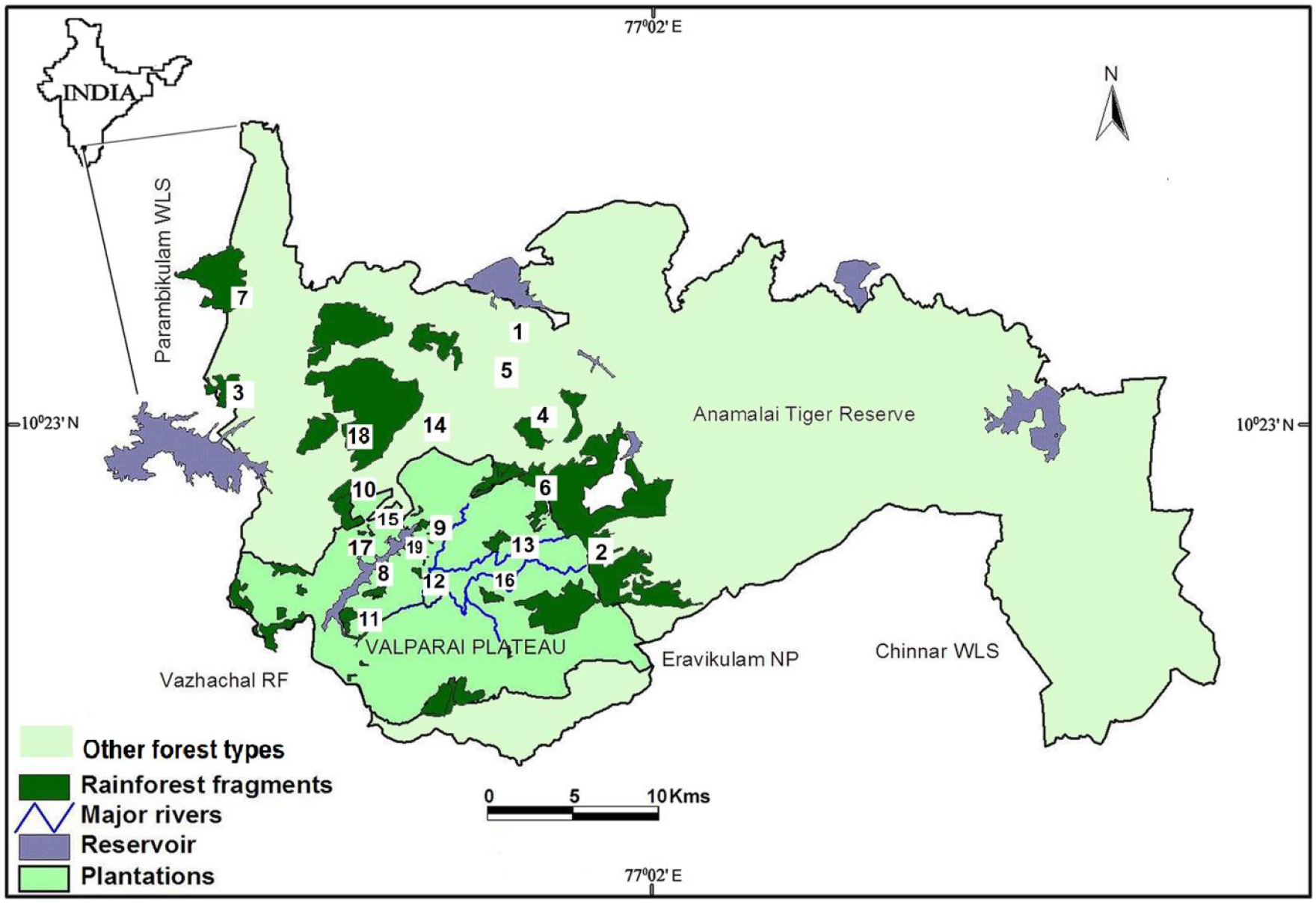
Map of Anamalai hills, Western Ghats, India with numbered study fragments. (1) Aliyar dam, 2) Akkamalai, 3) Anaikundi, 4) Andiparai, 5) Attakatty, 6) Iyerpadi 7) Karian_shola, 8) Korangumudi, 9) Monica_estate, 10) Monomboly, 11) Nirar_dam, 12) Pannimedu, 13) Puthuthottam, 14) Sethumadai 15) Shekkalmudi 16) Sirukundra 17) Uralikal, 18) Varagaliyar and 19) Varattuparai WLS: Wildlife sanctuary; RF: Reserve Forest; NP: National Park

### Host sampling

Between Oct 2013 and Oct 2015, faecal samples were collected from populations of mammalian wildlife. We collected fresh faecal samples, non-invasively during the day, on transects (400 m-3 km in length). For large and medium herbivores and primates, we followed individuals and collected fresh faeces when animals defaecated. For elusive species such as carnivores, we identified home range based on secondary information and faecal samples were identified based on morphology and also using nearby secondary signs such as pug-marks or hoof-prints. To avoid sampling the same individual repeatedly, only one sample of a host species was collected from each spot and the sample source was either marked or removed. To avoid contamination from soil, samples were collected from the inside of the bolus or only top pellet was collected from a heap. We immediately fixed each sample in 10% formaldehyde solution (50 ml), labelled the containers with the information of origin (fragment name, date, Time and host species) and stored them at room temperature until parasitological screening. Differences in sampling effort can confound the comparison of diversity among replicates. We accounted for differences in number of host species encountered by calculating richness estimates with the assumption that each faecal sample represents single individual. We used bootstrap, which is a resampling method for estimating the whole sampling distribution of richness by sampling with replacement from the original sample and can offer greater precision than jackknife estimates, especially when sample sizes are small ^61^.

### Parasite sampling

Employing both the flotation and sedimentation techniques (NaNO3 solution), we screened the faecal samples for the presence of helminth eggs, larvae and protozoan cysts ^62^. For each parasite concentration technique, we examined two slides per sample. Slides were examined under a light microscope (400X). Eggs and cysts were first examined at 10× magnification and then their size was measured with a micrometre eyepiece (0.1 μm) at 40× magnification. To facilitate identification of parasite eggs, we often added a drop of Lugol’s iodine solution to the slides, which highlighted detailed structures. In addition, photographs of each parasite species have been archived and are available for examination by request to the corresponding author. We identified parasites to the lowest possible taxonomic level using published keys ^63,64^. Differences in sampling effort can confound the comparison of diversity among replicates. We accounted for differences in number of parasite taxon encountered by calculating richness estimates with the assumption that each faecal sample represents single host individual. We used bootstrap, which is a resampling method for estimating the whole sampling distribution of richness by sampling with replacement from the original sample. Bootstrap can offer greater precision over jackknife estimator, especially when sample sizes are small ^61^. This method is particularly recommended for parasite richness estimation ^65^.

### Land use data

In Anamalai hills, land use change manifests in largely three forms—presence of human settlements, plantations and livestock foraging. There are only few large (>1000 ha) fragments that are legally protected and thus undisturbed. Many of the studied fragments share more than one type of land use change. For instance, some fragments with human settlements may also have livestock present. For the current study, we identified 18 fragments with land use change, out of which 15 (83.3%) had plantation, in contrast to three (16.7%). Eleven (61.1%) of the fragments have significant livestock foraging pressure, in contrast to seven (38.9%) fragments without livestock. Finally, ten (55.6%) of the fragments had human settlements within them, in contrast to eight (44.4%) without settlements.

### Habitat fragmentation data

To measure effect of habitat fragmentation, we used fragment size and isolation distance between fragments. According to the equilibrium theory of island biogeography, organism dispersal probability declines as distance between islands increases, reducing rates of immigration and, in turn, reducing diversity (MacArthur & Wilson, 1963, 1967; Whittaker & Fernandez-Palacios, 2007). Assuming each forest fragment as an island, their isolation was summarized with an isolation index which was calculated as the sum of the square root of the distances to the nearest equivalent (no smaller than 80% of size) or larger fragment (Dahl, 2004). Data on fragment size, distance between fragments and presence of human settlements, plantations and livestock were collected from earlier studies from Anamalai hills ^45,60^.

### Data analyses

To assess the effects of land use change and habitat fragmentations on bootstrap estimate of parasite taxon richness, we created two different linear mixed effects models ^66^. Each model included random effects of host species and fragments to account for multiple observations within each fragment (across host species) and across fragments. In the land use model, the predictor variables (fixed effects) were presence of plantation, human settlement and livestock. The predictor variables for the habitat fragmentation model were fragment size and fragment isolation index. In both the models, we incorporated both bootstrap estimates of host species richness and host body mass as co-predictors as these were known to effect parasite richness. We retrieved host body mass data from online ecological database ^67^. After fitting these model to the data, we also compared and selected the best fit model using lowest AIC value ^68^. At the end, diagnostics were run to check distribution of the residuals for each model. This analysis was conducted in the lme4 package ^69^. We also assessed the effects of land use change and habitat fragmentation on bootstrap estimates of host species richness using two linear models. In the land use model, the predictor variables were presence of plantation, human settlement and livestock. The predictor variables for the habitat fragmentation model were fragment size and fragment isolation index. We followed the same strategy as described above for model fitting, fitting diagnostics and model selection. Finally, we tested whether land use change homogenized the composition of the parasite community. We used a multivariate nonparametric Analysis of Variance (permAnoVa; 1,000 permutations) based on the Jaccard dissimilarity index for a matrix of parasite presence/absence. We calculated the variance of homogeneity of parasite communities within each fragment based on disturbed vs. undisturbed divisions using the betadisper function of the vegan package in R ^70^.

## RESULTS

### Sample diversity

From 19 forest fragments, we collected 4,056 mammalian faecal samples belonging to 23 mammalian wildlife species and two livestock species—domestic goats (*Capra aegagrus*) and cattle (*Bos taurus*). Analyses were done only on wildlife samples. Number of samples varied from 41 in Uralikal to 495 in Puthuthottam (Table 1). Number of samples for each host species varied between six in Otter (*Lutra lutra*) and 623 in gaur (*B. gaurus*). In total, seven protozoa (18.42%) and 32 helminth (81.58%) species were recorded, including five trematodes, five cestodes and 20 nematodes. At least seven different parasites, belonging to different parasite groups, were recorded in ≥20 different host species—protozoa *Coccidia sp.* (23 hosts); cestodes *Hymenolepis nana* (20 hosts) and *Moniezia sp.* (22 hosts); and nematodes *Gongylonema sp.* (20 hosts), *Strongyloides* sp. (23 hosts), *Trichuris sp.* (24 hosts) and *Ascaris sp.* (26 hosts). On the other hand, cestode *Dipylidium sp.* and nematode *Parascaris sp.* were found only in civet and Indian porcupine (*Hystrix indica*) samples, respectively.

**Table 1.** Bootstrap estimate of host richness in each fragment of Anamalai hills, India

| *Fragment name* | *Bootstrap estimate*<br>*of host species*<br>*richness* | *SE* | *Sample size* |
| --- | --- | --- | --- |
| Aliyar_dam | **21.9** | **0.8** | **210** |
| Akkamalai | **18.9** | **0.7** | **199** |
| Anaikundi | **20.1** | **0.8** | **150** |
| Andiparai | **22.6** | **1** | **256** |
| Attakatty | **11.7** | **0.6** | **15** |
| Iyerpadi | **25.5** | **1** | **398** |
| Karian_shola | **25.3** | **0.9** | **244** |
| Korangumudi | **24.9** | **0.8** | **356** |
| Monica_estate | **20.4** | **0.9** | **124** |
| Monomboly | **26.6** | **1** | **181** |
| Nirar_dam | **24.2** | **0.9** | **167** |
| Pannimedu | **17.5** | **1** | **55** |
| Puthuthottam | **24.9** | **0.8** | **426** |
| Sethumadai | **18.1** | **0.9** | **65** |
| Shekkalmudi | **16.0** | **0.8** | **42** |
| Sirukundra | **17.9** | **0.8** | **127** |
| Uralikal | **11.5** | **0.5** | **41** |
| Varagaliyar | **21.6** | **1** | **162** |
| Varattuparai | **23.4** | **0.9** | **397** |

### Host and parasite diversity and disturbance

For parasite diversity analysis the human disturbance model was the best fit (Table 2). Parasite diversity was significantly driven by presence of plantation (estimate = 4.779, CI_Profile_ = 0.326 – 9.232, t = 2.103, p<0.05). Presence of livestock had a substantial but not significant positive effect (estimate = 3.209, CI_Profile_ = −0.052 − 6.366, t = 1.992, p>0.05). Effects of settlement, host richness and host body mass on parasite richness were not significant (Figure 2). For host diversity analysis the human disturbance model was again the best fit (Table 3). Presence of plantation was the only predictor that had a significant positive effect on host diversity (estimate = 10.798, CI_Profile_ = 2.302 - 19.294, t = 2.726, p<0.05)—almost half of all host species occur in plantations. Although presence of livestock did not have a significant effect, its wide confidence interval was mostly on the positive side suggesting potential positive impact—limited by sample size—on host richness (estimate = 5.602, CI_Profile_ = −0.639 − 11.843, t = 1.925, p>0.05). Similarly, presence of human settlement did not significantly affect host richness, however, the substantial effect was mostly on the negative side, suggesting potential negative effect on host diversity (estimate = −4.112, CI_Profile_ = −10.717− 2.492, t = − 1.335, p>0.05).

**Table 2.** Comparison between two different models to explain bootstrap estimate of parasite taxon richness in Anamalai hills, India

| *Models* | *K* | *logLik* | *AIC* | *delta* | *weight* |
| --- | --- | --- | --- | --- | --- |
| *Plantation + Settlement + Livestock +*<br>*Host richness +Host body size* | 9 | -667.54 | 1353.07 | 0 | 0.887 |
| *Fragment size + Isolation index + Host*<br>*richness +Host body size* | 8 | -670.59 | 1357.19 | 4.112 | 0.113 |

**Table 3.** Comparison between two different models to explain bootstrap estimate of host species richness in Anamalai hills, India

| *Models* | *K* | *logLik* | *AIC* | *delta* | *weight* |
| --- | --- | --- | --- | --- | --- |
| *Plantation + Settlement + Livestock* | 5 | -46.73 | 103.47 | 0 | 0.971 |
| *Fragment size + Isolation index* | 4 | -51.25 | 110.5 | 7.029 | 0.029 |

**Figure 2.**
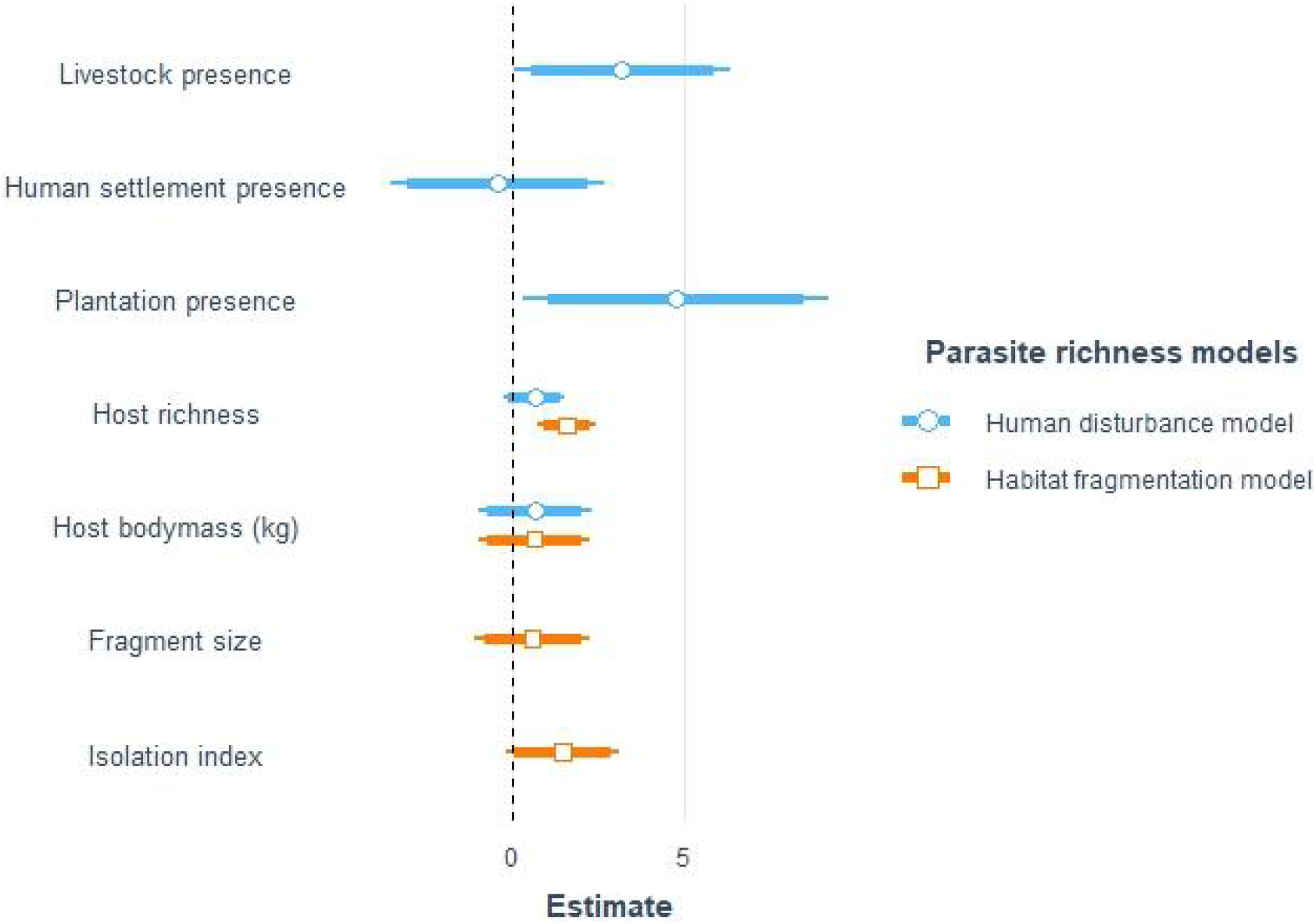
Unstandardized effect size of predictor variables on bootstrap estimate of parasite taxon richness in rainforest fragments of Anamalai hills, India. Estimates were plotted to scale. Intercepts were omitted to avoid distortion of scale. Disturbance model was the best fitted model based on AIC. Confidence intervals are represented by the lines around the points— thick (α = 0.10) and thin (α = 0.05).

We recorded 12 parasites (ten helminths and two protozoa) that occurred only in plantations. Six of the ten helminths were nematodes (60%), while rest were trematodes (30%) and one cestode (10%). Fragments without plantations did not harbour any parasite taxon exclusively, which means parasites in those undisturbed fragments also occured in plantations. Fragments with livestock harboured three parasite taxa (two nematodes and one cestode) exclusively relative to their undisturbed counterpart. However, only one parasite taxon (*Taenia sp.*) exclusively occurred in livestock disturbed fragments, while other two nematodes also occurred in the plantations. Its counterpart undisturbed fragments only harboured one taxon exclusively (*Paragonimus* sp.), which however also occurred in plantations. Finally, settlements harboured three nematode taxa exclusively in comparison to their undisturbed counterpart. Only one of these taxa (*Uncinaria sp.*) were exclusive to settlements across all fragments. Undisturbed counterpart of settlements harboured only one parasite taxon (*Sarcocystis sp.*) exclusively.

### Parasite and host homogeneity

Parasite communities within disturbed forest fragments were not significantly more homogeneous than the undisturbed ones due to presence of either plantations (F = 2.58, p>0.05), livestock (F = 0.04, p>0.05) or settlements (F = 3.55, p>0.05). Host communities within plantations (TukeyHSD; p<0.05; Figure 4a) and human settlements (TukeyHSD; p<0.05; Figure 4b) were, however, significantly more homogeneous than undisturbed fragments. Finally, we did not find any of the disturbance variables to significantly alter the parasite community composition between undisturbed and disturbed fragments.

## DISCUSSION

Our results reveal that rainforest fragments with plantations (and potentially with livestock) in Anamalai hills harbour significantly higher parasite diversity than undisturbed fragments. Interestingly, some of the modified fragments (at least, fragments with plantations) also has significantly more host diversity than the undisturbed fragments, however host diversity was not found to significantly affect parasite diversity.

In Anamalai hills, plantations (coffee, tea and cardamom) had more mammalian wildlife species richness than the undisturbed fragments. This was not particularly a surprising result because studies have reported similar high richness in vertebrate species from plantation within wildlife habitats ^71,72^. In fact, earlier studies from Western Ghats also found high vertebrate richness within or around plantations with large variations depending on plantation types, from open tea to more shaded coffee and cardamom plantations ^30,73,74^. The reason for such increased host diversity is thought to be an increase in habitat heterogeneity within plantations. Increased habitat heterogeneity is thought to generate greater diversity of niches consequently facilitating cooccurrence of many species ^75,76^. However, such increase in species richness is often accompanied by more generalist and wide-ranging species being more abundant within the plantations and a loss of community heterogeneity relative to undisturbed habitats ^77–79^. We found similar loss of heterogeneity for host species in disturbed habitats with plantations and settlements (Figure 4).

Effect of livestock presence on host species richness was positive but not statistically significant at α = 0.05. The effect, however, was significant at α = 0.10, which suggested potential, but weak effect, which was reflected by the almost equal number of wildlife species recorded from these two groups of fragments (n_Livestock_ = 20 and n_Undisturbed_ = 22). Interaction between livestock and wildlife is complicated. For instance, while a number of studies found evidence of competitive exclusion between livestock and large herbivore ^80^, may other recorded resource sharing between these two groups ^81,82^. Yet still, may other studies did not find any relationship between the two ^83^. The outcome of the interaction may depend on the ecological similarity between the two groups (Niche overlap), availability of natural resources that may vary between habitats (between low to high productivity) and also degree of behavioural habituation by the wildlife. The wildlife community that we studied was an ecologically broad one consisting of wildlife with very different ecology. Therefore, while some of the species—such as spotted deer and sambar deer, who were found only in the undisturbed fragments—may face resource competition from livestock grazing, others (for example, small carnivores and primates) may not face any competition. In addition, many large herbivores, such as gaur and elephants, who despite resource competition may still use the disturbed fragments as corridors contributing to host richness. These processes together may explain almost similar host species richness between fragments with and without livestock grazing.

We did not find any significant effect of human settlement on host diversity but the trend is negative (Figure 3). While human settlement may attract and facilitate generalist and weedy species with high tolerance for disturbance (for example, rodents, which were not sampled in the present study), many elusive species such as carnivores may be adversely affected and may prefer to avoid fragments with settlements ^84^. Still, we recorded overall a large host species richness (host richness_Settlement_ = 19, host richness_Undisturbed_ = 23) from around the settlement in Anamalai hills. This could be explained by the facts that many of these settlements may attract wildlife with unintentionally supplemented resources such as planted fruit trees ^60^. Additionally, the high level of fragmentation of the landscape meant large herbivores and carnivores may not have much choice but to disperse through human settlements ^54,56^. We did not find evidence of habitat fragmentation (fragment size and isolation) influencing host species richness in Anamalai hills (Figure 3). This is in line with findings from across studies that effects of fragmentation on species communities are often weak ^85^. Effects of habitat fragmentation on species diversity is highly context-specific and varies considerably between animal groups, ecosystems and kinds of human activities prevalent in the landscape ^85–89^. In Anamalai hills, habitat fragmentation is widespread, which likely disrupt animal movement to some extent but, in the absence of hunting, perhaps not substantially. For instance, studies recorded use of certain plantation as corridors to connect with isolated undisturbed habits ^51,52,74^. However, the adverse outcome of these movement through human-dominated habitats is the increase in wildlife-human conflict ^54,56^.

**Figure 3.**
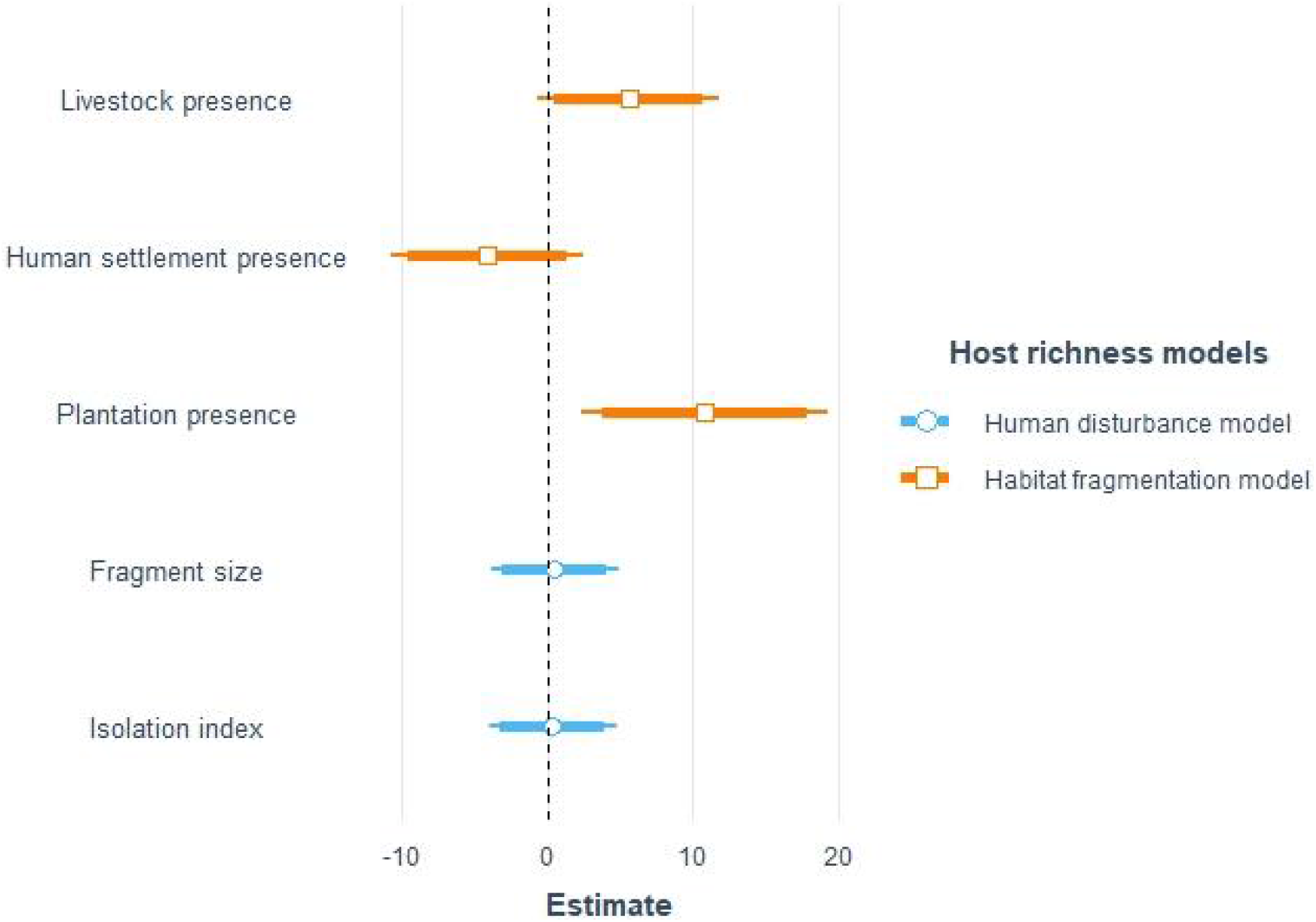
Unstandardized effect size of predictor variables on bootstrap estimate of host species richness in rainforest fragments of Anamalai hills, India. Estimates were plotted to scale. Intercepts were omitted to avoid distortion of scale. Disturbance model was the best fitted model based on AIC. Confidence intervals are represented by the lines around the points— thick (α = 0.10) and thin (α = 0.05). Host sample sizes are given in Table 1.

**Figure 4.**
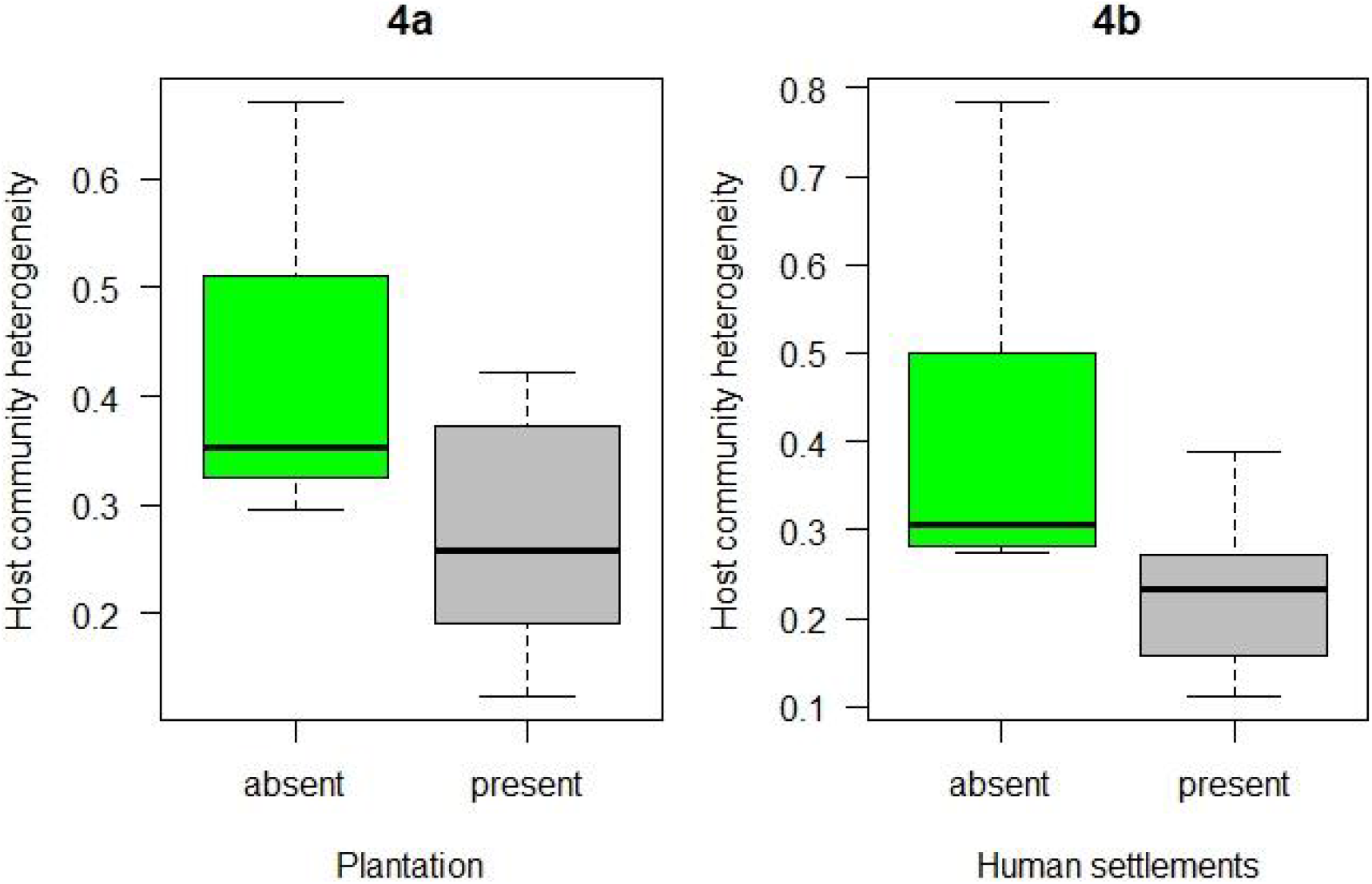
Host community heterogeneity between undisturbed (absent) and disturbed (present) rainforest fragments of Anamalai hills, India. Community heterogeneity is the within group dispersion values based on Jaccard distance for presence/absence data.

Among the different types of land use change in Anamalai hills, plantations had the strongest positive effect on parasite diversity (Figure. 2). Increase in number of parasite taxa in modified fragments ranged between one to ten, with eight parasite taxa that were recorded exclusively in these fragments (Table 4). However, this increased richness in disturbed fragments were likely not driven by host richness as host richness had a small and statistically not significant effect on parasite richness in the disturbance model (Figure 2). This is in contrast to the predominant patterns across most studies on parasite diversity that found host richness to be the strongest predictor of parasite richness ^19–22^. However, there could be potential deviations from this rule, particularly due to human impacts ^21,90,91^. For instance, many human parasites may spillover to wildlife (anthropozoonoses) as humans regularly come in contact with wildlife ^92–97^. Humans may also introduce many non-wildlife species such as feral dogs, cats in addition to livestock into wildlife habitats and these species may share parasites with wildlife ^26^. In such cases, parasite richness in wildlife would be more than in the undisturbed fragments. Indeed, all but one (*Schistosoma sp.*) of the parasites that we found exclusively in plantations also occurred in cattle (Table 4). Surprisingly, wildlife parasite taxa that were present in the livestock foraging fragments did not occur in cattle samples from the same fragments. This was also the case for the wildlife parasites that only occurred in settlement but not in undisturbed fragments. We did not find any significant effect of host body mass on parasite diversity (Figure 2). This is in contrast many studies that found a significant relationship between these two variables ^98,99^. On the other hand, many other empirical studies that did not find any relationship between body mass and parasite richness when accounting for host phylogenetic relationships ^100,101^. Such contradictory results may suggest that relationship between host body mass and parasite diversity is a factor of body mass and life history traits, which vary between ecologically different groups of hosts ^22^. Thus, the broad ecological diversity among host species in the present study might have confounded this relationship.

**Table 4.** Parasite taxa that were found only in disturbed or undisturbed fragments in Anamalai hills, India. Highlighted parasite taxa were found in the corresponding fragment group exclusively. 1. Current study; 2. Natural History Museum parasite database, London, UK

| Parasite taxa | Parasite group | Family | In livestock samples<sup>1</sup> | Known human case<sup>2</sup> |
| --- | --- | --- | --- | --- |
| **Plantation only** |  |  |  |  |
| **Baylisascaris sp.** | **Nematodes** | **Ascaridoidea** | **Present** | **present** |
| **Nematodirus sp.** | **Nematodes** | **Trichostrongyloidea** | **Present** | **present** |
| **Enterobius sp.** | **Nematodes** | **Oxyuroidea** | **Present** | **present** |
| Dictyocaulus sp. | Nematodes | Trichostrongyloidea | Absent | absent |
| Uncinaria sp. | Nematodes | Ancylostomatoidea | Absent | present |
| **Schistosoma sp.** | **Trematodes** | **Schistosomatidae** | **Absent** | **present** |
| Metastrongylus sp. | Nematodes | Metastrongyloidea | Absent | present |
| **Clonorchis sp.** | **Trematodes** | **Opisthorchiidae** | **Present** | **present** |
| **Toxoplasma sp.** | **Apicomplexa** |  | **Present** | **present** |
| **Isospora sp.** | **Apicomplexa** |  | **Present** | **present** |
| Paragonimus sp. | Trematodes | Paragonimidae | Present | present |
| **Dipylidium sp.** | **Cestodes** | **Dilepididae** | **Present** | **present** |
| **Livestock presence only** |  |  |  |  |
| Dictyocaulus sp. | Nematodes | Trichostrongyloidea | Absent | absent |
| **Taenia sp.** | **Cestodes** | **Taeniidae** | **Absent** | **present** |
| Metastrongylus sp. | Nematodes | Metastrongyloidea | Absent | present |
| **Undisturbed (Livestock) only** |  |  |  |  |
| Paragonimus | Trematodes | Paragonimidae | Present | present |
| **Settlement only** |  |  |  |  |
| Dictyocaulus sp. | Nematodes | Trichostrongyloidea | Absent | absent |
| Uncinaria sp. | Nematodes | Ancylostomatoidea | Absent | present |
| Metastrongylus sp. | Nematodes | Metastrongyloidea | Absent | present |
| **Undisturbed (Settlement) only** |  |  |  |  |
| **Sarcocystis sp.** | **Apicomplexa** |  | **Present** | **present** |

Our results did not find any significant effect of habitat fragmentation on parasite richness (Figure 2). This was expected as we did not find any effect of fragmentation on host richness either. This lack of relationship between fragmentation and parasite diversity could also be an outcome of large home ranges and low habitat specialisation of most of the host species in our study. Many of the species that we sampled were large herbivores or carnivores (e.g., *Elephas maximus*, *Bos gaurus*, *Panthera tigris*, *Panthera pardus*) with lang home range and they disperse across fragments. The level of fragment isolation (Median distance = 30.2 km) may not be a deterrent to their dispersal. Similarly, many host species in the study community such as *Macaca radiata*, *Sus scrofa*, *Viverricula indica*, are habitat generalists. According to the distribution-abundance relationship hypothesis ^6^, smaller, fragmented habitats may be conducive for these generalist wildlife with high reproductive rates. These hosts may then spread parasites across habitats, independent of the level of habitat fragmentation.

## CONCLUSION

With data on 40 parasites from 23 host species collected from 19 rainforest fragments with different types of land use change, we demonstrated that land use changes increased parasite diversity and presence of potential spillover of parasites from livestock to wildlife. We also showed that the observed pattern of parasite diversity was not driven by habitat fragmentation.

One of the limitations of this study was that it could not test the effect of land use change and habitat fragmentation on the relationship between host density and parasite diversity. Host density is an important predictor of parasite diversity and in nature, host density is linked to host ecology (e.g., home range). However, land use change can unpredictably change host density, which may have a complex outcome for parasite diversity. It will thus be worthwhile in future to explore this question in the present system. Additionally, with the present evidence of potential anthropozonosis, it will be important in future to compare parasite from the present study to samples from humans, livestock and commensal animals in the fragments. Finally, as far as land use change and habitat fragmentation of wildlife habitats in India are concerned, the present study represents a case study with particular relevance for tropical rainforest habitats. However, there exists a large diversity in habitats and levels of disturbance in India. Given the increased threat to wildlife health from anthropogenic environmental change, it will thus be crucial for wildlife conservation to study the patterns of parasite diversity in other types of habitats, especially those with already threatened wildlife.

## Data availability

The datasets generated and analysed during the current study are available from the corresponding author on request.

## Acknowledgements

DC gratefully acknowledges Steffen Foerster for initial discussions on model selection. DC also used part of Fulbright fellowship to work on the current paper, for which he gratefully acknowledges support of US-India Education Foundation (US-IEF), New Delhi. GU gratefully acknowledges funding awarded to him by Department of Biotechnology, Government of India (Ref. No. BTPR4503/BCE/8/899/2012 dated 09-11- 2012).

## Author Contributions

Sampling and data collection were done by DMR, ST and DC. DC conducted data analysis and wrote manuscript. GU conceived, designed and lead the study, organised funding and reviewed the manuscript.

